# A single decreasing ramp friction sprint for torque-cadence relationship assessment during cycling

**DOI:** 10.1101/2025.02.11.637613

**Authors:** Pablo Rozier-Delgado, Maximilien Bowen, Marion Dussauge, Pierre Samozino, Baptiste Morel

## Abstract

This study aimed to introduce and validate a novel method for assessing dynamic fatigue components through a single-sprint test, addressing the limitations of traditional multi-sprint evaluations. We tested this method on twenty-one participants by computing torque-cadence relationships from two iso-friction sprints at varying friction levels (3% and 9% of body mass), the traditional combination of these iso-friction sprints and a novel decreasing ramp friction sprint (Fr_D_). The accuracy of this new method through fatigue was also tested with ten 6-s Fr_D_ sprints interspersed with a 24-s passive rest. Fr_D_ outperformed single iso-friction sprints and provided accurate and valid torque-cadence relationship’s parameters estimates (T_0_, C_0_, and P_max_) with systematic bias < 3%, typical error of estimate < 6% and very high r^2^ (median of 0.962). The quality of the input data from this method was also high, as evidenced by the well-distributed and wide-range cadence spectrum (51.3% of C_0_; skewness = −0.51, p < 0.05) and was maintained throughout the fatiguing exercise. Our novel method not only allows the dynamic fatigue components evaluation with only one sprint but also maintains accuracy and validity across varying fatigue states, offering significant advantages for both research and practical applications.

## Introduction

Neuromuscular fatigue can be defined as the inability to maintain the required or expected force output (Bigland-Ritchie and Woods, 1984). Disregarding the fatiguing exercise, whether it is continuous, intermittent, mono-articular or locomotion tasks (implying various contraction modes), isometric maximal voluntary contraction (IMVC) force is the most popular criterion of fatigue (Brownstein et al., 2021), as it represents an experimentally convenient measurement and allows for fatigue aetiology investigation (when combined with evoked stimulation; Krüger et al. 2018). Yet, measuring only the isometric force may not represent all aspects of neuromuscular fatigue. Indeed, as fatigue occurs within a task, dynamic output (e.g. power) can decrease, while there is no change in IMVC output (Krüger et al., 2019), leading to crucial fatigue manifestation omission.

One more comprehensive way to assess force production is through the force-velocity relationship, which allows for continuous analysis of the capabilities throughout the entire contraction velocity spectrum. Over the years, the force-velocity relationship (Eq. 1) has been well studied from contractile properties in isolated muscles (Hill, 1938) to multi-joint tasks, such as cycling, jumping, sprint running, and bench pressing (Jiménez-Reyes et al., 2014; Morin et al., 2010; Rahmani et al., 2017; Vandewalle et al., 1987). During cycling, this relationship can be expressed as the torque-cadence relationship *T* (*C*) defined by (i) *T*_0_ (in N · m), the theoretical torque production corresponding to a null cadence; (ii) *C*_0_ (in rpm), the theoretical maximal cadence until which torque can be produced (Dorel, 2018a).

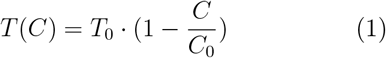

Since the first torque-cadence test reported in scientific literature on a cycle ergometer (Dickinson, 1928), methods have changed and improved due to technological progress and innovative protocols that allow shorter procedures (Driss and Vandewalle, 2013; Vandewalle and Driss, 2015). In 1979, Pirnay and Crielaard (1979) proposed a protocol of eight to ten sprints performed against different loads to obtain maximum cadence for each load. This method was then improved and the amount of loads was reduced (two loads used as warm-up and familiarisation and four to five different loads) due to the linearity of the torque-cadence relationship (Vandewalle et al., 1985, 1987). Still, this large number of sprints is time-consuming as it needs sufficient recovery time between sprints and, therefore, does not allow quick torque-cadence profile evaluation. Yet, assessing dynamic force production capacity within a short delay can be crucial when evaluating fatigue at several time points over or after the exercise. Therefore, to overcome this limitation, it is possible to compute the average torque and velocity at each downstroke during the acceleration phase to obtain multiple torque-cadence data points within a single sprint (Arsac et al., 1996). Although this method is much quicker, the velocity range covered by experimental points is quite small during iso-inertial sprint against one given load. Consequently, to guarantee accurate estimation of *C*_0_ and *T*_0_, protocols still require multiple sprints with different friction to cover a sufficiently large spectrum of velocities (Dorel et al., 2005; Dorel, 2018b).

On this basis, Krüger et al. (2020) proposed a biomechanical model to determine the optimal load that permits, regardless of an individual’s fitness level, to better distribute the data around maximal power. This approach utilises a single sprint that maps cadences on both sides of the power-cadence relationship allowing better estimations of *T*_0_ and *C*_0_ parameters. However, this method still involves constant friction load and inertia as the only ways of varying cadences. Therefore, despite this better cadence distribution, the optimal load method covers a similar cadence spectrum than the traditional method (only 34% of *C*_0_) and still has non-homogeneous data distribution along this spectrum, compromising the accuracy of the estimation of the parameters (i.e., *T*_0_ and *C*_0_).

Still, some studies have investigated the effects of intermittent or continuous exercise on fatigue by using single iso-friction sprints (Bogdanis et al., 2007; Buttelli et al., 1996). For example, Hautier et al. (1998) observed a greater decrease in *C*_0_ than *T*_0_ in repeated sprint exercise. Similar results were found with repeated sprint exercise during running, with a greater decrease in maximal theoretical velocity (Jiménez-reyes et al., 2018). However, it is worth noting that during an iso-inertial running sprint, force capacities at low velocities are tested in the very first seconds of the test. In contrast, force capacities at high velocities are tested after 5 or 6s, where fatigue can already appear. These methodological aspects may affect fatigue-induced alteration in the force-velocity relationship, with a more pronounced effect at high velocities. That is why, dynamic force capacities testing should be short (<3 or 4 s) to reduce the fatigue induced by the test (Dorel, 2018a). Considering these temporal constraints (i.e., a single sprint lasting less than 3-4s), the mechanical conditions of traditional cycle ergometers (inertial wheel, fixed friction force) and interindividual variability do not allow a protocol to sweep across a wide range of cadences in one single sprint without additional fatigue caused by the test itself. In order to test dynamic force capacities in the shortest time and over the widest spectrum of velocities, one single iso-inertial sprint against a decreasing friction could be interesting to consider. This study aimed to propose and test the validity of a new method to assess the cycling torque-cadence relationship. This method, a single sprint with decreasing friction load, could be efficiently implemented in fatigue protocols while still allowing for a high-quality torque-cadence assessment (i.e., with sufficient data points, well spread across an important cadence spectrum and with high coefficients of determination). It was hypothesised that this method allows for (i) a greater cadence spectrum covered and less skewed cadence data compared to the single friction load sprint method and (ii) a valid estimation of the torque-cadence relationship parameters compared to the gold standard method (multiple sprints with constant friction method) and (iii) can be implemented in protocols of fatigue with maintained assessment quality.

## Methods

### Participants

Twenty-one participants (age: 24.1 ± 6.7 years old; height: 178.5 ± 5.5 cm; body mass: 73.3 ± 6.8 kg) volunteered for this study. All participants were free from pain or injury and provided written informed consent prior to starting the study. This study was conducted in compliance with the Declaration of Helsinki and was approved by the local Human Research Committee.

### Methodology

Each volunteer participated in a unique experimental session comprising (i) a warm-up, (ii) multiple torque-cadence evaluations with fixed and ramp friction conditions, and (iii) a repeated sprint exercise. First, a 9-minute standardized warm-up was performed before the test with 5 min of cycling at 1 W · kg^−1^, 3 min at 2 W · kg^−1^, and 1 min at 3 · W kg^−1^. After two familiarization sprints, the participants performed three 6-second sprints in a randomised order with 3 minutes of passive recovery. Two sprints were performed with an iso-friction load of 3% and 9% of body mass (Fr_I3_ and Fr_I9_, respectively), and a third sprint was performed with a decreasing ramp friction (Fr_D_) from 1 to 0 N · kg^−1^ in 6 s. Finally, repeated sprint exercise (ten 6-s Fr_D_ sprints interspersed with a 24-s passive rest) was conducted. Before each sprint, the participants were instructed to remain seated on the saddle and to produce the highest acceleration possible during each pedal cycle, while the experimenters provided vigorous encouragement. All participants used toe clips.

### Material (*Apparatus*)

A Monark cycle ergometer (Monark LC6, Stockholm, Sweden) was instrumented with both a strain gauge with its amplifier (Futek FSH04207; 444.822 N; M6×1; 500Hz Gain 1.9N/V) that measures the frictional force applied by the belt to the flywheel and with an optical encoder (Hengstler 600 points per revolution) that assesses the linear displacement of a virtual point on the flywheel periphery. The friction force applied to the belt was controlled using a motorised linear actuator. The external torque (in N · m) and cadence (in rpm) produced by the subject were calculated using the gear ratio (52/14), the radius of the wheel (0.257 m) and considering the torque to overcome the flywheel inertia (I=1.01 kg · m^2^). Torque and cadence signals were sampled (200 Hz) and stored in a custom Labview program (National Instruments Corporation, Austin, TX) via a 16-bit analogue-to-digital interface card Ni DAQ (NI-USB6210 National Instruments Corporation, Austin, TX).

### Data analysis

#### Processing

To remove frequencies that are non-pedalling high-frequency phenomena (e.g., bike vibrations, electromagnetic noise) and to preserve the pedalling signal lower than 300 rpm (≡ 5 Hz), cadence and torque data were filtered with a zero-phase Butterworth low-pass filter (3^rd^ order, cut-off frequency: 5 Hz) and averaged for each pedal downstroke of the acceleration phase (Samozino et al., 2007). A threshold of 95% of the maximal cadence was used to consider only the acceleration phase and remove the cadence plateau. For all participants, the first downstroke was incomplete and, as a result, was systematically deleted. The Eq. 1 was fitted using the non-linear least squares method with the remaining data of Fr_I3_, Fr_I9_, Fr_D_, and by combining Fr_I3_ and Fr_I9_ data (Fr_IC_, method considered here as the gold standard method). Any residual absolute *Z*-score greater than 3 was considered an outlier and was removed before a second fitting procedure. From this final fitting, *T*_0_, *C*_0_, the optimal cadence (C_opt_ in rpm; Eq. 2) and the maximal power output (*P*_*max*_ in W; Eq. 3) were computed for each sprint method (Vandewalle et al., 1987). For each profile, as a characterisation of input data quality, the number of points (n_points_), the coefficient of determination (r^2^), the 95% confidence interval of the parameters, the cadence spectrum covered (C_spectrum_; Eq. 4) as well as the skewness, an indicator of distribution asymmetry (the closer to 0, the more symmetrical) were computed. All subsequent data processing was performed using MATLAB (R2021b, MathWorks, USA).

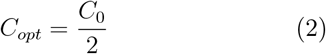

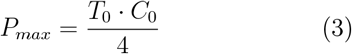

#### Statistical analysis

For both initial capacity evaluation and repeated sprint exercise, the assumption of normality on *T*_0_, *C*_0_, *P*_*max*_, and input data quality parameters (n_points_, r^2^, spectrum, and skewness) was checked using the Shapiro–Wilk test and evaluated graphically using QQ plots. Any residual absolute *Z*-score greater than 3 on *T*_0_ and *C*_0_ was considered an outlier, and the entire participant was then removed from the analysis. Then, to test the concurrent validity of Fr_D_ compared to Fr_IC_, correlation and Bland-Altman analyses were performed for *T*_0_, *C*_0_, and *P*_*max*_. The coefficient of correlation (Pearson’s r), mean bias and typical error of estimate (expressed in raw values and as a coefficient of variation in %) were computed (Hopkins et al., 2009). Sphericity was checked through the Mauchly test, and the Greenhouse-Geisser transformation was applied when necessary. Then, a one-way analysis of variance (ANOVA) for repeated measures was conducted between the methods (Fr_I3_, Fr_I9_, Fr_IC_, and Fr_D_) on *T*_0_, *C*_0_, *P*_*max*_, and parameters describing the input data quality (i.e., the cadence spectrum (Eq. 4) and skewness) with Bonferroni corrected post-hoc tests if a significant effect was reported. When the assumption of normality was not confirmed (i.e., n_points_ and r^2^), a Friedman test was conducted instead with Conover post-hoc tests when appropriate.

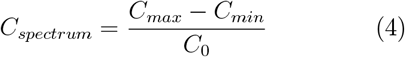

With C_max_ and C_min_ being respectively the maximal and minimal cadence measured during the sprint.

Statistical significance was set at p-value < 0.05. Partial eta squared 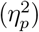 was reported as an estimate of effect size for ANOVA, with 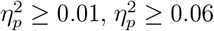, and 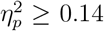 used as small, moderate, and large effects, respectively (Cohen, 1988).

Finally, a paired comparison between first and last sprint was conducted to test the fatigue effect during repeated sprint exercise on the torque-cadence relationship parameters (*T*_0_, *C*_0_, and *P*_*max*_) and the input data quality parameters (n_points_, r^2^, C_spectrum_, and skewness). A Student paired t-test was performed between the first and last sprint in the repeated sprint exercise when the normality assumption was checked; otherwise, the Wilcoxon test was performed. Cohen’s d or Cliff’s *δ* (Cliff, 1993) were reported as estimates of the effect size for parametric and non-parametric tests, respectively. Cohen’s d threshold values of 0.2, 0.6, 1.2, 2.0 and 4.0 were used to represent small, moderate, large, very large, and extremely large effects, respectively (Hopkins et al., 2009), whereas Cliff’s *δ* threshold values of 0.11, 0.28 and 0.43 were used to represent small, medium, and large effects, respectively (Vargha and Delaney, 2000). Statistical analysis was performed using JASP (version 0.17.1, University of Amsterdam, Netherlands) and MATLAB (R2021b, MathWorks, USA). Results are displayed as mean ± s.d. unless otherwise specified.

## Results

### Input data quality

The number of points (i.e., pedal downstroke), coefficient of determination (r^2^), range of covered spectrum (Eq. 4), and the skewness for each method are presented in table 1. There was a statistically significant difference in the number of points depending on which sprint method was used (*χ*^2^(3) = 51.2, p<0.001). Both Fr_I3_ and Fr_I9_ had a similar number of data points covering the torque-cadence relationship (12(3) vs. 12(2) as median(IQR), p-value = 0.235). However, Fr_IC_ had significantly more points on the torque-cadence relationship compared to Fr_D_ (Hodges-Lehmann estimate = +9.5 [8;11], Cliff’s *δ* = 0.98, p-value = 0.003) The last point to be taken into account for the *T* (*C*) relationship for Fr_I3_, Fr_I9_, Fr_IC_ and Fr_D_ occurred at respectively 4.7, 5.7, 5.8 and 5.2 seconds, in average.

**Table 1.**
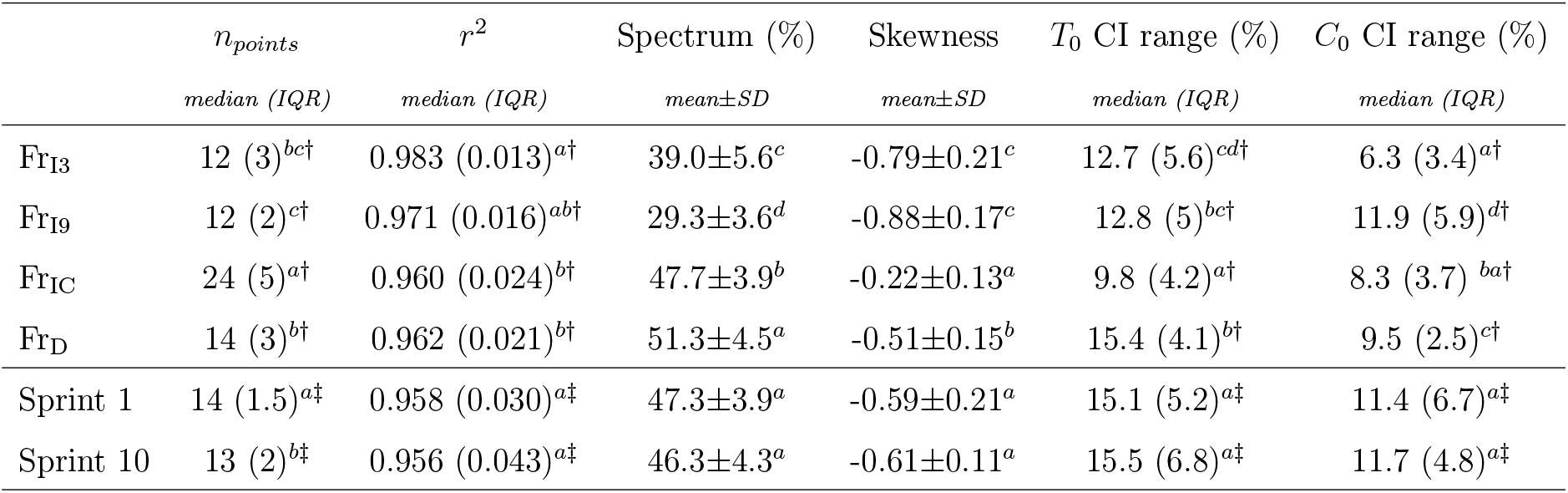
Parameters of input data quality for each method and between first and last sprints in repeated sprint exercise. The number of points (n_points_), coefficients of determination (r^2^), skewness, and 95% confidence interval ranges (*T*_0_ and *C*_0_ CI range) are displayed. Values with different superscript letters in a column are significantly different from each other (p-value < 0.05). †: Non-parametric Friedman test used with Conover post-hoc analysis. ‡: Non parametric Wilcoxon test

### Validity

The results of the analysis of the concurrent validity are shown in table 2. Pearson correlations, absolute bias and limits of agreement from the Bland-Altman analysis are reported in Fig. 2.

**Table 2.**
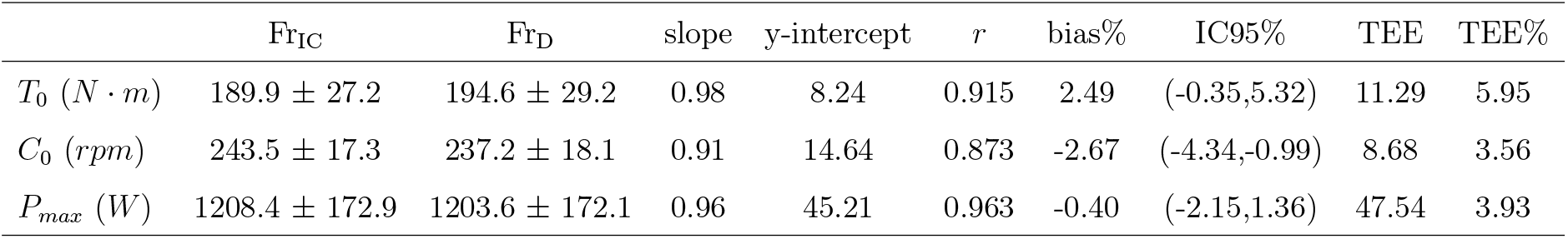
Difference between Fr_D_ and Fr_IC_ method with their associated reliability statistics.

| | Fr<sub>IC</sub> | Fr<sub>D</sub> | slope | y-intercept | $r$ | bias% | IC95% | TEE | TEE% |
| --- | --- | --- | --- | --- | --- | --- | --- | --- | --- |
| $T_0$ (N · m) | 189.9 ± 27.2 | 194.6 ± 29.2 | 0.98 | 8.24 | 0.915 | 2.49 | (-0.35,5.32) | 11.29 | 5.95 |
| $C_0$ (rpm) | 243.5 ± 17.3 | 237.2 ± 18.1 | 0.91 | 14.64 | 0.873 | -2.67 | (-4.34,-0.99) | 8.68 | 3.56 |
| $P_{\text{max}}$ (W) | 1208.4 ± 172.9 | 1203.6 ± 172.1 | 0.96 | 45.21 | 0.963 | -0.40 | (-2.15,1.36) | 47.54 | 3.93 |

There was a statistically significant difference in the coefficient of determination depending on which sprint method was used (*χ*^2^(3) = 17.3, p-value <0.001). The r^2^ values for Fr_I3_, Fr_I9_, Fr_IC_, and Fr_D_ were similar (table 1). However, Fr_I3_ had a significantly higher r^2^ compared to Fr_IC_ and Fr_D_ (Hodges-Lehmann estimate [CI95%] = +0.018 [0.006;0.030], Cliff’s *δ* = 0.58, p=0.05 and +0.018 [0.009;0.033], Cliff’s *δ* = 0.58, p-value <0.001, respectively) and Fr_I9_ had a significantly higher r^2^ compared to Fr_D_ (Hodges-Lehmann estimate [CI95%] = +0.008 [0.001;0.016], p-value = 0.048, Cliff’s *δ* = 0.28) but lower compared to Fr_I3_ (Hodges-Lehmann estimate [CI95%] = −0.012 [−0.021;-0.001], p-value = 0.048, Cliff’s *δ* = 0.37). There was a statistically significant difference in cadence spectrum covered depending on which sprint method was used (*F*_3,60_ = 149.8, p-value <0.001, 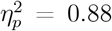), with Fr_D_ covering a larger range of spectrum than Fr_I3_ and Fr_I9_ (diff = +12.3% [+9.3; +15.4], d = 1.93 and diff = +22.0% [+18.9; +25.1], d = 5.99 respectively, all p-values < 0.001) and even covering a larger spectrum than Fr_IC_ (diff = + 3.6% [+0.5; +6.7], d = 0.79, p-value = 0.013). Finally, there was a statistically significant difference in skewness depending on which sprint method was used (*F*_3,60_ = 88.9, p-value <0.001, 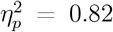). Post-hoc analysis showed no significant difference in skewness between Fr_I3_ and Fr_I9_ (diff = 0.09 [−0.03;0.21], p-value = 0.289). However, Fr_D_ was significantly less skewed than Fr_I3_ and Fr_I9_ (diff = +0.28 [0.16; 0.41], d = 1.24, p-value<0.001 and diff = +0.37 [0.25; 0.50], d = 1.79, p-value <0.001, respectively; table 1), with Fr_IC_ being significantly less skewed than Fr_D_ (diff = 0.29 [0.17; 0.41], d = 1.42, p-value < 0.001).

### Repeated sprint exercise

Figure 3 shows the data related to the evolution of the relative covered cadence spectrum before and after fatigue. The parameters related to input data quality are presented in table 1. Two outliers were detected and removed from the analysis (participant 11 with repeated sprint 1 and participant 20 with repeated sprint 10). All the measured torque production capacity variables exhibited a significant decrease between the first and last sprint. *T*_0_ decreased significantly from 190.4 ± 30.0 N · m to 173.9 ± 25.3 N · m (difference: −16.6 [−24.2; −8.9] N · m, t(18) = 4.5, p-value < 0.001, d = 1.04 [0.47; 1.59]), *C*_0_ decreased significantly from 229.5 ± 15.3 rpm to 215.4 ± 21.0, rpm (difference = −14.1 [−22.9; −5.4] rpm, t(18) = 3.4, p-value = 0.003, d = 0.78 [0.26; 1.29]), and *P*_*max*_ decreased significantly from 1140.0 ± 164.2 W to 978.6 ± 155.8 W (difference = −161.4 [−223.0; −99.8] W, t(18) = 5.5, p-value < 0.001, d = 1.26 [0.65; 1.86]). Despite fatigue, the relative covered cadence spectrum did not change significantly (difference = −1.0% [−3.1; 1.1], t(18) = 1.02, p-value = 0.319), and neither did r^2^ (Hodges-Lehmann = −0.005 [−0.026; 0.013], W = 82, p-value = 0.623), and the skewness (difference: −0.026 [−0.126; 0.075], t(18) = 0.54, p-value = 0.596) (table 1). However, number of data points decreased significantly (Hodges-Lehmann estimate: −1.5 [−2.5; 0], W = 79, Cliff’s *δ* = 0.42, p-value = 0.019).

## Discussion

The purpose of this study was to propose and test the validity of a single iso-inertial sprint with decreasing friction to assess the cycling torque-cadence relationship. The main results are that decreasing ramp friction (Fr_D_) sprint i) allows valid assessment of *T*_0_, *C*_0_, and *P*_*max*_; ii) covers a much greater range of cadences than iso-friction loads, with similar r^2^ values; iii) In a fatigued state, the Fr_D_ sprint still covers a higher spectrum of cadences, r^2^, and better data spreading (skewness) compared to iso-friction load sprints, allowing for qualitative assessment of dynamic force capabilities for fatigue investigation.

First, the values obtained for *T*_0_, *C*_0_, and *P*_*max*_ align well with the previous *T* (*C*) relationship evaluation for a healthy active population. In this study, average *P*_*max*_ was 1208 ± 173 W (16.5 W · kg^−1^), average *T*_0_ was 190 ± 27 N · m and *C*_0_ was 243.7 ± 17.3 rpm, in accordance with Dorel et al. (2010) who found 1132 W (16.2 W · kg^−1^), 181 N · m and 236 rpm. Similarly, other studies investigating young healthy populations found values ranging around 14 W · kg^−1^ for relative *P*_*max*_, between 150 to 198 N · m for absolute *T*_0_ and between 236 to 240 rpm for absolute *C*_0_ (Hautier et al., 1996; Kordi et al., 2019). When comparing the proposed Fr_D_ method to the gold standard multi-sprint Fr_IC_, extremely low bias and typical error of estimates (respectively under 3% and 7%) were evidenced. Systematic bias of 4.7 N · m and −6.3 rpm were significant for *T*_0_ and *C*_0_, respectively, but associated with Cohen’s D of 0.35 and −0.16 showing, at worst, a small effect.

As for the different indicators of input data quality (i.e., spectrum, skewness), Krüger et al. (2020) observed a ≈ 30% cadence spectrum coverage with the traditional intermediate iso-friction load sprint (7.5% of body mass). Our study did not investigate this specific friction load but tested single load conditions around this value. We found corresponding values, with Fr_I3_ and Fr_I9_ ranging between 39% and 29% (in average) of the covered cadence spectrum. Conversely, with only one sprint, the Fr_D_ covers more than 50% of the cadence spectrum, higher than Fr_IC_. Moreover, single iso-friction sprints had cadences more skewed towards lower values (−0.79 and −0.88 for Fr_I3_ and Fr_I9_ respectively) than Fr_D_ sprints (−0.51). Negative skewness could deteriorate the *T*_0_ estimation during the fitting as more data is present at high cadences. Although Fr_IC_ had the best skewness (−0.22), a single Fr_D_ sprint allows for an important improvement in skewness compared with iso-friction loads. Finally, in addition to having a broad spectrum and well-distributed data, the Fr_D_ method also features a spectrum that is similarly centered around C_opt_ (50% of *C*_0_) compared to Fr_IC_ (see Fig. 1). Krüger et al. (2020) also reported r^2^ = 0.89 ± 0.06 for iso-friction sprints at 7.5% of body mass load. In this study, we reported r^2^ > 0.96 for all traditional iso-friction-loaded sprints (Fr_I3_, Fr_I9_, and Fr_IC_). The higher r^2^ can be explained by the different processing methods used for the *T* (*C*) relationship computation. In this study, outlier data points were retrieved after the first fitting as some data points can be submaximal (Rudsits et al., 2018). This combined better-distributed data, with a large range of cadences covered and high r^2^, allows for a better mathematical extrapolation of *T*_0_ and *C*_0_ (Krüger et al., 2020) and a confident assessment of torque-cadence relationships.

**Figure 1.**
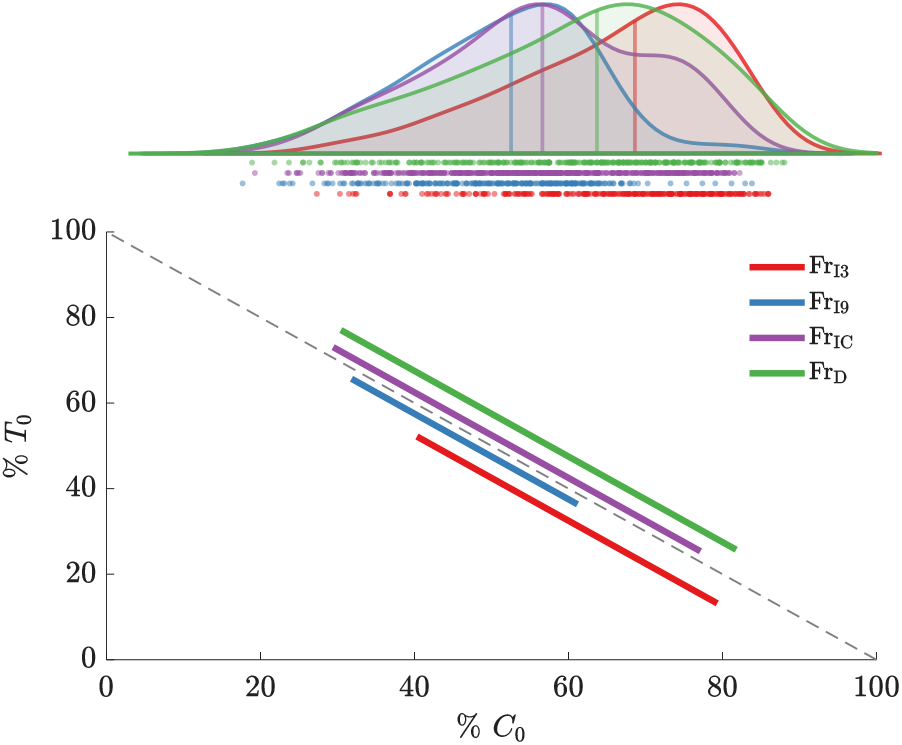
Overall distribution and cadence spectrum covered for initial evaluation. Iso-friction sprint at a 3% (Fr_I3_) and 9% (Fr_I9_) of body mass loads are displayed in red and blue, respectively, whereas the combination of both iso-friction sprint (Fr_IC_) and decreasing ramp friction sprint (Fr_D_) are displayed in purple and green, respectively.

**Figure 2.**
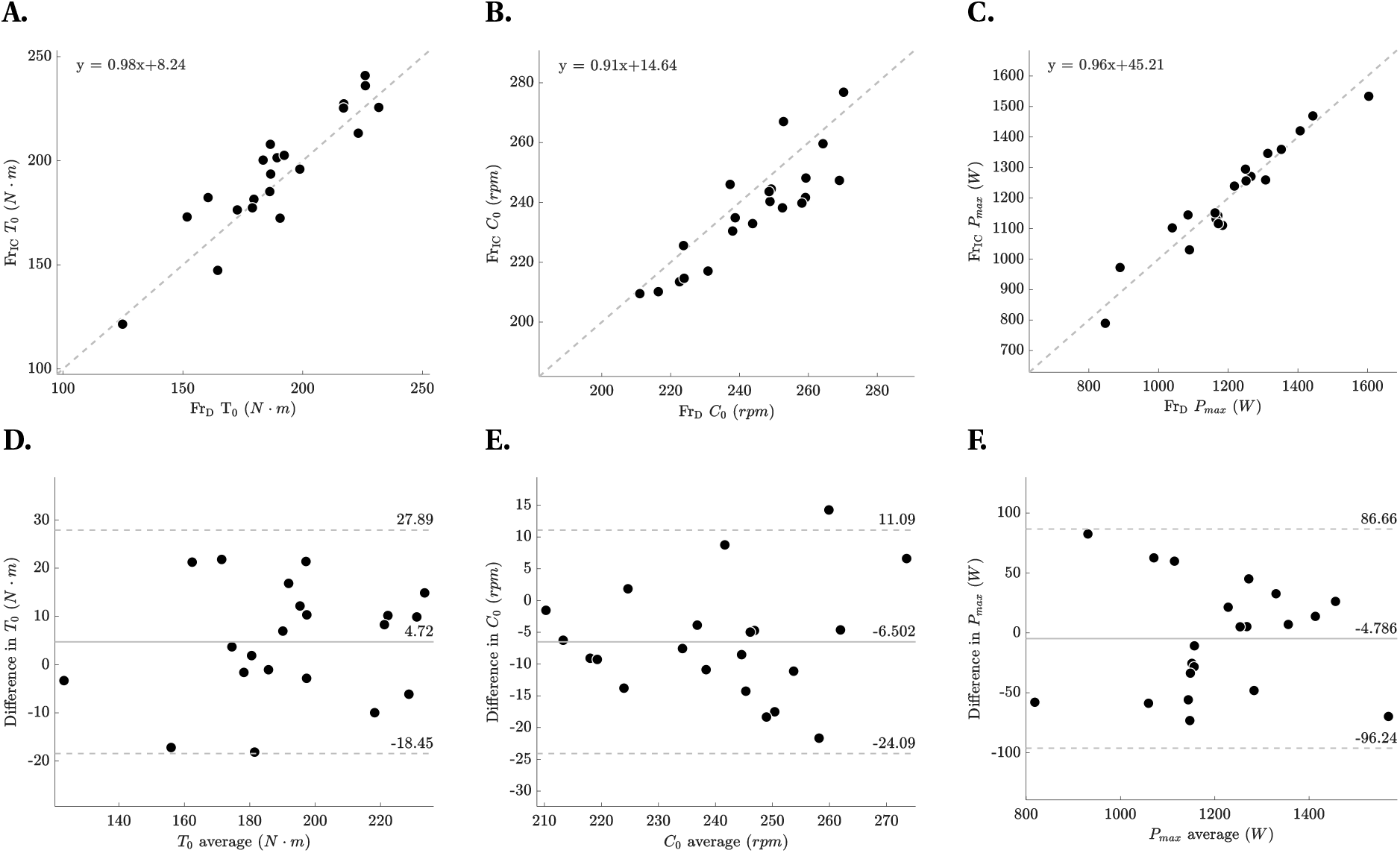
Validity between Fr_IC_ and Fr_D_ method. Linear relationship for **(A)** *T*_0_ **(B)** *C*_0_ **(C)** *P*_*max*_ and Bland-Altman plot depicting levels of agreement for **(D)** *T*_0_, **(E)** *C*_0_ and **(F)** *P*_*max*_, including absolute bias estimate (grey line) and both lower and upper 95% limits of agreement (grey dashed lines).

**Figure 3.**
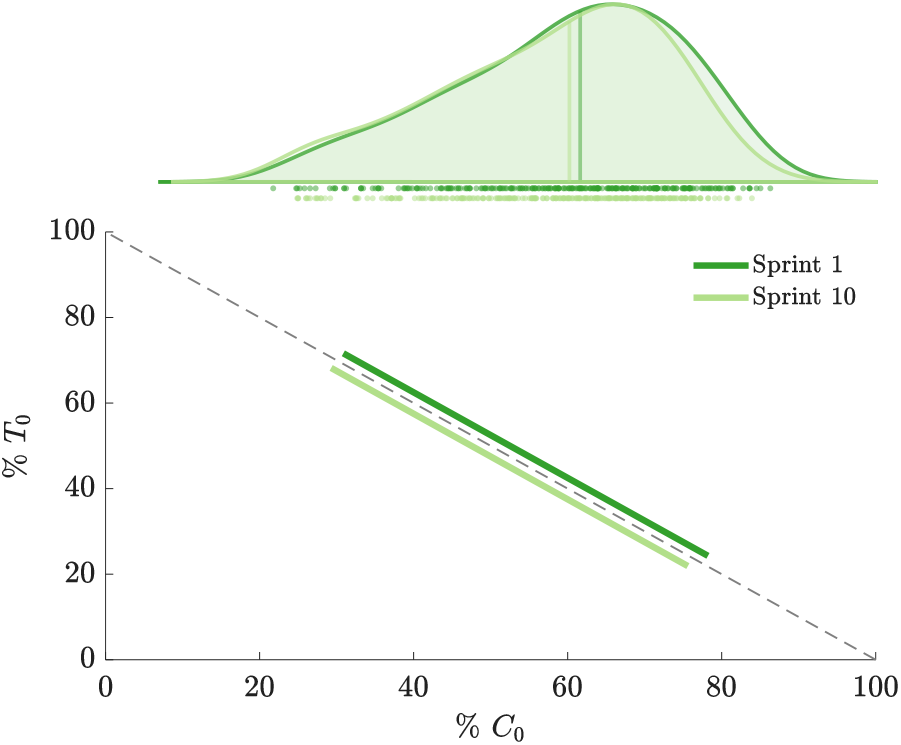
Overall distribution and cadence spectrum covered during repeated sprint protocol. Decreasing ramp friction sprint (Fr_D_) for the first and last sprint of the repeated sprint exercise are displayed in dark green and light green, respectively.

**Figure 4.**
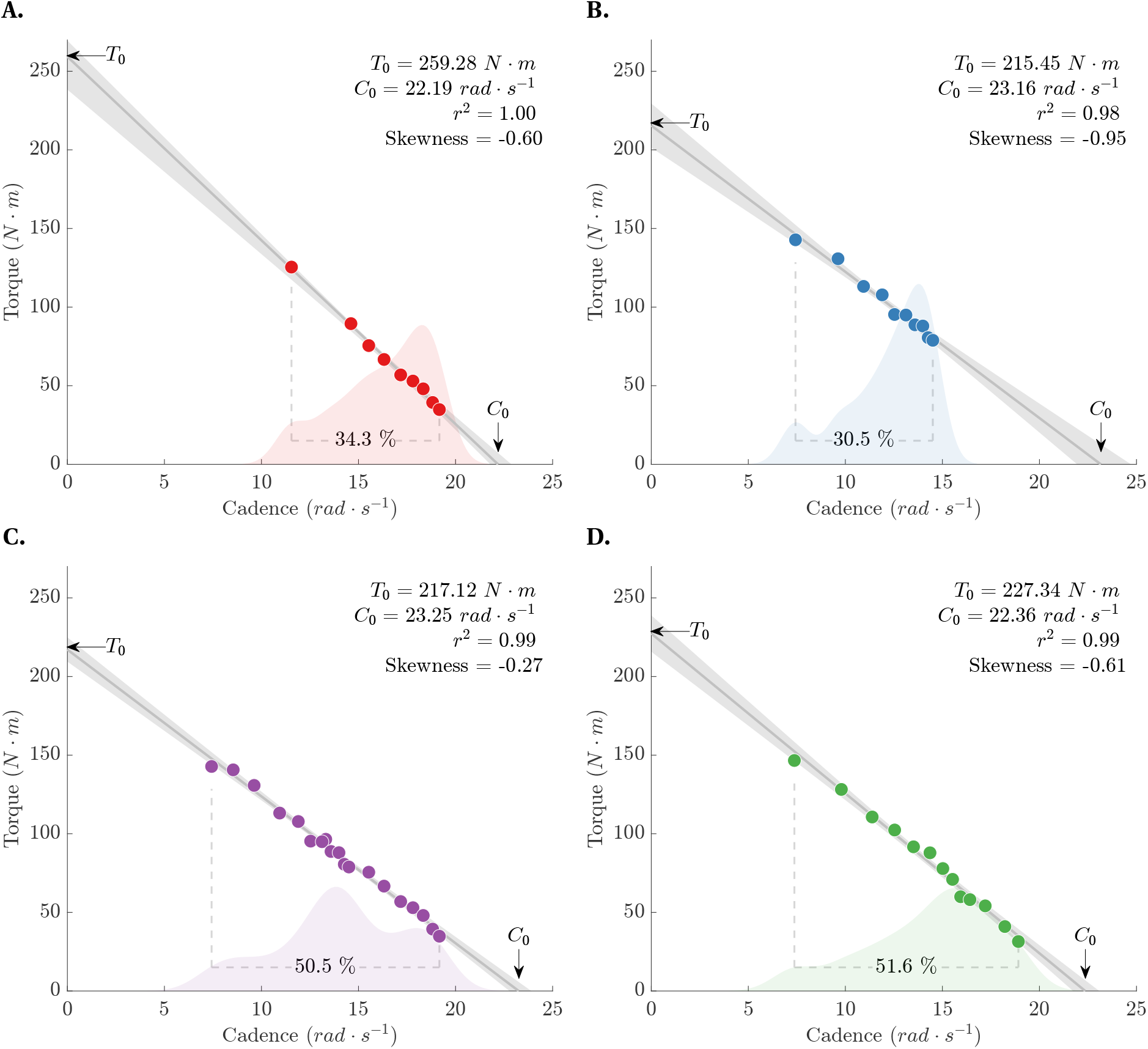
Typical torque-cadence relationships for one participant. **(A)** Fr_I3_ method; a sprint with iso-friction load fixed at 3% of body mass **(B)** Fr_I9_ method; a sprint with iso-friction load fixed at 9% of body mass **(C)** Fr_IC_ method; the combination of both 3% and 9% body mass iso-friction sprints and **(D)** Fr_D_ method; a sprint where friction decreases from 1 to 0 N · kg^−1^ in 6 seconds. The grey line represents the torque-cadence profile with its associated 95% confidence interval (grey patch). Parameters estimate, coefficient of determination and skewness are displayed on the top-right while the cadence spectrum and the data distribution are displayed below the data points.

Similarly, all the input data quality indicators were superior to the previously proposed optimal load iso-friction sprint (Fr_OPT_; Krüger et al. 2020). The Fr_D_ sprint presented higher r^2^ (0.96 (0.03) for Fr_D_ VS 0.92 ± 0.06 for Fr_OPT_). This higher r^2^ value can be explained by the same reason mentioned *supra* (i.e., varying processing methods). In addition, Krüger et al. (2020) reported a ≈ 35% range of the spectrum covered by the optimal load method. While this method is easy to implement, as no specific friction assessment is required and represents an improvement compared to traditional iso-friction sprints, it still covers a relatively low cadence spectrum. Yet, this spectrum is important for accurate relationships, as a low spectrum can lead to large systematic and random errors for both *T*_0_ and *C*_0_ parameters estimation.

The major advantage of Fr_D_ method over the classic assessment of *T* (*C*) relationship is when performed in a fatigued state because whether it is the traditional or the optimal load method, the constant friction applied to the belt is chosen and optimised for a non-fatigued state. As soon as fatigue occurs, there is an important loss of cadence capacity (Hautier et al., 1998) that could prevent the participant from reaching *P*_*max*_, as was the case for three individuals in the Krüger et al. (2020) study. With important cadence left-skewed data and a low spectrum of cadences covered (i.e., ≈ 25%), large errors can appear in the *T* (*C*) parameter extrapolation. In this study, we found very high input data quality parameters, even during a fatiguing task. With a ≈ 161 W loss and despite a 14.1 rpm loss for *C*_0_ and a lower number of points, there was no difference with pre-fatigue measurements on r^2^, covered spectrum, and skewness. Therefore the Fr_D_ method allows for a very high-quality *T* (*C*) relationship assessment, even in a fatigued state.

It should be noted that the repeated sprint exercise induced a 14% loss in *P*_*max*_, whereas Krüger et al. (2020) reported a 27% loss in *P*_*max*_. Higher fatigue shall lead to a higher decrease in reached cadences, and further studies should investigate whether it reduces the quality of the assessment or not. Nevertheless, in theory, friction evolving to a zero value at the end of the sprint should make it possible to maintain a very large covered range, whatever the magnitude of fatigue. Another limitation is that the Fr_D_ was still more skewed to the left than the Fr_IC_. Indeed, the multiple sprints traditionally performed to assess the *T* (*C*) relationship (e.g. Linossier et al., 1996) allows well-distributed data around *P*_*max*_. However, Fr_D_ is still a very high-quality assessment method and allows quick *T* (*C*) evaluations with a sufficiently high r^2^ and range of cadences to be covered. Furthermore, the method presented here proposes a linear decrease in friction. Nothing would prevent adjusting the friction decay according to another function in order to homogenise the distribution of points over the spectrum covered.

As recently demonstrated, fatigue can alter static or dynamic capacities differently (Krüger et al., 2018, 2019). Even dynamic contractions appear to be specifically sensitive to fatigue, depending on the speed condition tested (Morel et al., 2019, 2015). Thus, the ability to quickly (≈5s) assess the maximal theoretical torque capacity (*T*_0_) and maximal theoretical cadence (*C*_0_) capacity represents a major asset for fatigue evaluation in both continuous and intermittent exercises. Whether it is possible to increase the slope of the force applied to the belt to scan, in just a few seconds, a large spectrum of cadences is yet to be investigated.

In conclusion, a robust *T* (*C*) relationship assessment on a stationary cycle ergometer often requires several sprints with a large spectrum of cadences and a very high r^2^, so that *T*_0_ and *C*_0_ are well extrapolated from the fit. The use of an instrumented ergometer to control the force applied to the belt can permit the assessment of this *T* (*C*) relationship within only one sprint with a conserved quality and improved cadence spectrum. As fatigue occurs, this new method offers an even bigger advantage, as the quality of the relationship is conserved regardless of the neuromuscular performance loss.

## Abbreviations

BM: Body mass
*T*_0_: Maximal theoretical torque
*C*_0_: Maximal theoretical cadence
*P*_*max*_: Maximal theoretical power
IMVC: Isometric maximal voluntary contraction
rpm: Revolutions per minute
Fr_D_: Decreasing ramp friction condition
Fr_IC_: Iso-friction combined condition

## Acknowledgments

Support was provided by the Agence Nationale de la Recherche [ANR-22-CE14-0073] to B. Morel with additional support to P. Rozier-Delgado [ANR-22-EXES-0017]. This work is part of the Force-Velocity-Endurance (FoVE) framework.

